# Long-term immune changes after COVID-19 and the effect of BCG vaccination and latent infections on disease severity

**DOI:** 10.1101/2024.12.16.628601

**Authors:** Kamila Bendíčková, Ioanna Papatheodorou, Gabriela Blažková, Martin Helán, Michaela Haláková, Petr Bednář, Erin Spearing, Lucie Obermannová, Julie Štíchová, Monika Dvořáková Heroldová, Tomáš Tomáš, Roman Panovský, Vladimír Šrámek, Marco De Zuani, Marcela Vlková, Daniel Růžek, Marcela Hortová-Kohoutková, Jan Frič

## Abstract

**Background:** Several years after the COVID-19 pandemic, the role of trained immunity in COVID-19 remains controversial, and questions regarding the long-term effects of COVID-19 on immune cells remain unresolved. We investigated the roles of Bacillus Calmette–Guérin (BCG) vaccination and latent infections in the progression of COVID-19 and sepsis.

**Methods:** We conducted a prospective analysis of 97 individuals recovering from mild-to-critical COVID-19 and 64 sepsis patients. Immune cell frequencies, expression of functional markers, and plasma titres of anti-*Toxoplasma gondii*/cytomegalovirus/BCG antibodies were assessed and their impact on disease severity and outcomes were determined. To examine monocyte responses to secondary challenge, monocytes isolated from COVID-19 convalescent patients, BCG vaccinated and unvaccinated volunteers were stimulated with SARS-CoV-2 and LPS.

**Results:** Post COVID-19 patients showed immune dysregulation regardless of disease severity characterized mainly by altered expression of activation and functional markers in myeloid (CD39, CD64, CD85d, CD11b) and lymphoid cells (CD39, CD57, TIGIT). Strikingly, post-critical COVID-19 patients showed elevated expression of CD57 in CD8^+^ T cells compared to other severity groups. Additionally, a higher frequency of CMV and *T. gondii* seropositive-alongside a lower frequency of BCG seropositive-patients were associated with severe and critical COVID-19. However, the monocyte response to stimulation was unaffected by the severity of COVID-19.

**Conclusion:** These findings highlight the long-term alterations of immune cells in post-COVID-19 patients emphasizing the substantial impact of COVID-19 on immune function. However, our data showed no relationship between previous BCG vaccination and protection against SARS-CoV-2 infection.

## INTRODUCTION

The emergence of severe acute respiratory syndrome coronavirus 2 (SARS-CoV-2) and the subsequent global pandemic affected millions of people. While most cases of COVID-19 were mild, some escalated into severe complications, such as pneumonia. Recent research shows that SARS-CoV-2 can also cause long-lasting changes to the immune system [1, 2], potentially providing an immunological basis for the long-term effects of COVID-19. However, the long-term impact of COVID-19 severity on the immune system remains largely unknown.

The innate immune response is essential for ensuring the immediate defence against pathogens in a non-specific manner. The key cellular players include monocytes/macrophages, granulocytes and natural killer (NK) cells, which are the first cells to respond to pathogenic infection. Exposure to live pathogens or microbial molecules triggers innate immune cells, providing them with long-term hyper-responsiveness against non-specific antigens. This process is driven by profound cellular and functional reprogramming to elicit “trained immunity” (TI) [3, 4]. Characteristic example of this is the Bacillus Calmette–Guérin (BCG) vaccine, primarily developed against tuberculosis, that also provides off-target heterologous protection against non-mycobacterial infections[5, 6] as a result of innate immune reprogramming [3, 4]. Similarly, latent infections such as *Toxoplasma gondii* (*T. gondii)* and cytomegalovirus (CMV) were associated with an improved response of innate immune cells to unspecific pathogens in animal models [7, 8] as well as in primary human monocytes[7, 9]. While the molecular mechanisms are still being delineated [10], we know that TI is generally underpinned by the enhanced production of inflammatory cytokines (including TNF-α, IL-6, IL-1β) following re-exposure to unrelated pathogens [11, 12].

The clinical impact of TI is profound in the context of patients with a compromised host defence, such as those with post-sepsis immune paralysis, cancer, autoimmune diseases and inflammatory disorders. In the latter, exaggerated TI can contribute to disease pathogenesis [4, 13, 14]. Numerous studies have made associations between BCG vaccination status and COVID-19 outcomes, but no consensus has emerged. For example, several early epidemiological studies found that inhabitants of countries lacking a BCG vaccination policy were more prone SARS-CoV-2 infection and succumbing to COVID-19 [15, 16]. Although these studies suggested a potential protective effect of BCG vaccination, they cannot establish definitive causality due to several inherent biases [17]. Meanwhile, several randomised clinical trials evaluating single-dose or multiple-dose of BCG vaccination reported either protective effects [18-22] or no effect [23-28] on COVID-19 outcomes in high-risk patients. Finally, a post-hoc analysis of several recent studies found a beneficial effect of BCG vaccination on the overall survival of COVID-19 patients [29], with the emphasis on COVID-19 severity rather than incidence of infection [30].

Aside from COVID-19, the impact of BCG vaccination has also been considered in the context of other infections and even sepsis. Data suggests that the BCG vaccine elicits protection mainly against respiratory infections in elderly [31] and neonatal sepsis in newborns of vaccinated mothers [31-33]. Similarly, the contribution of latent infections in the protection against infection and sepsis progression has been studied. Nevertheless, the outcomes are again mixed: Latent CMV infection provided protection against bacterial pathogens *Listeria monocytogenes* and *Yersinia pestis* in a mouse-model[8], and latent *T. gondii* infection had a long-term impact on the phenotype and responsiveness to *T. gondii* re-infection of primary human monocytes[7], which may have important implications for innate immune responses to unrelated pathogens. On the other hand, mice chronically infected with *T. gondii* were more susceptible to sepsis induced by caecal ligation and puncture, and patients with sepsis and *T. gondii* seropositivity have a high mortality rate [9]. Therefore, further research is required to rule out the beneficial or deleterious role of latent infections in the outcome of sepsis or other infections.

To determine the long-term impact of COVID-19 on the immune cell frequencies and functionality, we established a prospective study, recruiting individuals who had recovered from COVID-19 with different severity and performed deep immunophenotyping. Since there is growing evidence for the role of TI in the ability to improve defence against viral respiratory infections [34, 35], we were interested in whether previous BCG vaccination or latent infections can provide protection against COVID-19 and impact sepsis survivorship. We also assessed the association between prior BCG vaccination and COVID-19/sepsis progression to determine whether there indeed exists a relationship between BCG vaccination status, latent infections and COVID-19/sepsis outcomes, including pneumonia severity and sepsis survivorship. We then analysed isolated monocytes from COVID-19 patients and BCG-vaccinated/non-vaccinated volunteers and tested their responses to lipopolysaccharide (LPS) and SARS-CoV-2 stimulation to investigate if COVID-19 or previous BCG vaccination induce enhanced pro-inflammatory responses in monocytes.

## METHODS

### Cohort demographics

The post-COVID-19 patient cohort comprised 104 adults with mild to critical COVID-19 who were recruited between October 2021 and September 2024. Seven post-COVID-19 patients were excluded from the analysis due to incomplete data. Disease severity classification followed the WHO guidelines [36]. Immunophenotyping of whole blood samples was performed on 78 patients. The cohort demographic characteristics are summarised in Table 1 and a comparison of the clinical characteristics of patients with severe and critical COVID-19 is shown in Supplementary Table 1. In total 64 patients with sepsis were recruited to the study between January 2023 and September 2024 (Table 2).

**Table 1:**
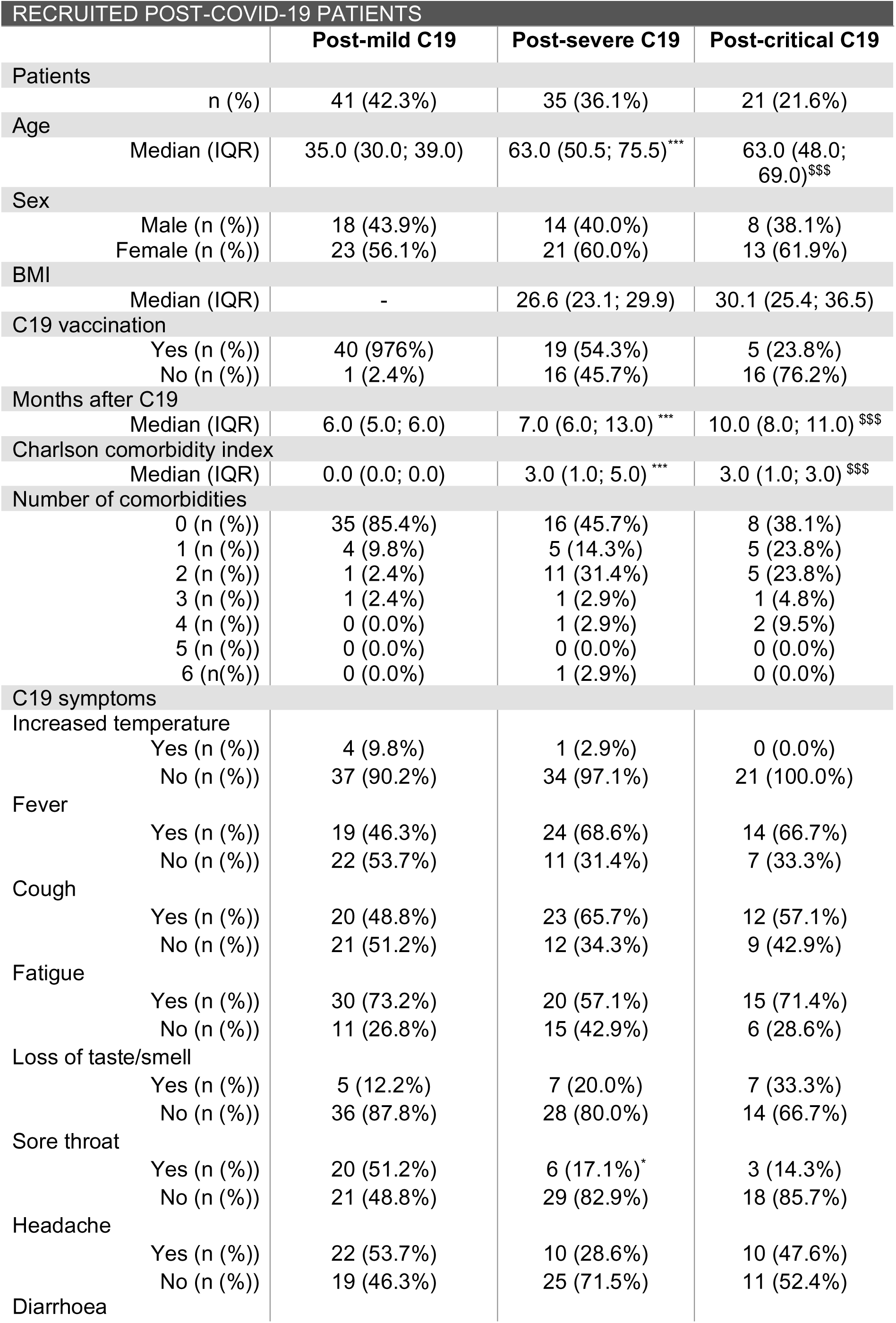
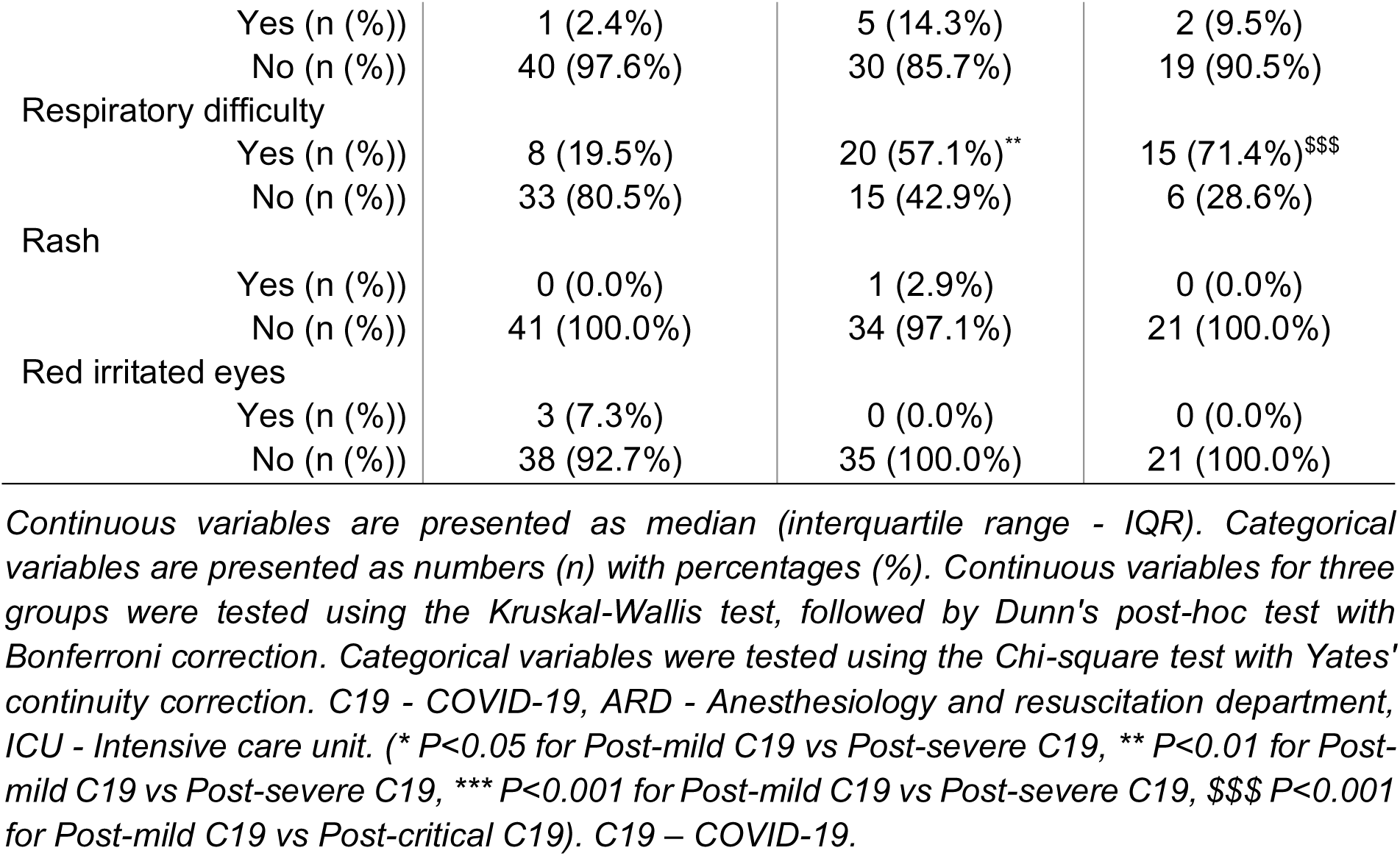
Demographic characteristics of post-COVID-19 cohort.

| RECRUITED POST-COVID-19 PATIENTS |  |  |  |
| --- | --- | --- | --- |
|  | Post-mild C19 | Post-severe C19 | Post-critical C19 |
| Patients |  |  |  |
| n (%) | 41 (42.3%) | 35 (36.1%) | 21 (21.6%) |
| Age |  |  |  |
| Median (IQR) | 35.0 (30.0; 39.0) | 63.0 (50.5; 75.5)*** | 63.0 (48.0; 69.0)\$\$\$ |
| Sex |  |  |  |
| Male (n (%)) | 18 (43.9%) | 14 (40.0%) | 8 (38.1%) |
| Female (n (%)) | 23 (56.1%) | 21 (60.0%) | 13 (61.9%) |
| BMI |  |  |  |
| Median (IQR) | - | 26.6 (23.1; 29.9) | 30.1 (25.4; 36.5) |
| C19 vaccination |  |  |  |
| Yes (n (%)) | 40 (97.6%) | 19 (54.3%) | 5 (23.8%) |
| No (n (%)) | 1 (2.4%) | 16 (45.7%) | 16 (76.2%) |
| Months after C19 |  |  |  |
| Median (IQR) | 6.0 (5.0; 6.0) | 7.0 (6.0; 13.0)*** | 10.0 (8.0; 11.0)\$\$\$ |
| Charlson comorbidity index |  |  |  |
| Median (IQR) | 0.0 (0.0; 0.0) | 3.0 (1.0; 5.0)*** | 3.0 (1.0; 3.0)\$\$\$ |
| Number of comorbidities |  |  |  |
| 0 (n (%)) | 35 (85.4%) | 16 (45.7%) | 8 (38.1%) |
| 1 (n (%)) | 4 (9.8%) | 5 (14.3%) | 5 (23.8%) |
| 2 (n (%)) | 1 (2.4%) | 11 (31.4%) | 5 (23.8%) |
| 3 (n (%)) | 1 (2.4%) | 1 (2.9%) | 1 (4.8%) |
| 4 (n (%)) | 0 (0.0%) | 1 (2.9%) | 2 (9.5%) |
| 5 (n (%)) | 0 (0.0%) | 0 (0.0%) | 0 (0.0%) |
| 6 (n (%)) | 0 (0.0%) | 1 (2.9%) | 0 (0.0%) |
| C19 symptoms |  |  |  |
| Increased temperature |  |  |  |
| Yes (n (%)) | 4 (9.8%) | 1 (2.9%) | 0 (0.0%) |
| No (n (%)) | 37 (90.2%) | 34 (97.1%) | 21 (100.0%) |
| Fever |  |  |  |
| Yes (n (%)) | 19 (46.3%) | 24 (68.6%) | 14 (66.7%) |
| No (n (%)) | 22 (53.7%) | 11 (31.4%) | 7 (33.3%) |
| Cough |  |  |  |
| Yes (n (%)) | 20 (48.8%) | 23 (65.7%) | 12 (57.1%) |
| No (n (%)) | 21 (51.2%) | 12 (34.3%) | 9 (42.9%) |
| Fatigue |  |  |  |
| Yes (n (%)) | 30 (73.2%) | 20 (57.1%) | 15 (71.4%) |
| No (n (%)) | 11 (26.8%) | 15 (42.9%) | 6 (28.6%) |
| Loss of taste/smell |  |  |  |
| Yes (n (%)) | 5 (12.2%) | 7 (20.0%) | 7 (33.3%) |
| No (n (%)) | 36 (87.8%) | 28 (80.0%) | 14 (66.7%) |
| Sore throat |  |  |  |
| Yes (n (%)) | 20 (51.2%) | 6 (17.1%)* | 3 (14.3%) |
| No (n (%)) | 21 (48.8%) | 29 (82.9%) | 18 (85.7%) |
| Headache |  |  |  |
| Yes (n (%)) | 22 (53.7%) | 10 (28.6%) | 10 (47.6%) |
| No (n (%)) | 19 (46.3%) | 25 (71.5%) | 11 (52.4%) |
| Diarrhoea |  |  |  |

|  |  |  |  |  |
| --- | --- | --- | --- | --- |
|  | Yes (n (%)) | 1 (2.4%) | 5 (14.3%) | 2 (9.5%) |
|  | No (n (%)) | 40 (97.6%) | 30 (85.7%) | 19 (90.5%) |
| Respiratory difficulty | Yes (n (%)) | 8 (19.5%) | 20 (57.1%)** | 15 (71.4%)\$\$\$ |
|  | No (n (%)) | 33 (80.5%) | 15 (42.9%) | 6 (28.6%) |
| Rash | Yes (n (%)) | 0 (0.0%) | 1 (2.9%) | 0 (0.0%) |
|  | No (n (%)) | 41 (100.0%) | 34 (97.1%) | 21 (100.0%) |
| Red irritated eyes | Yes (n (%)) | 3 (7.3%) | 0 (0.0%) | 0 (0.0%) |
|  | No (n (%)) | 38 (92.7%) | 35 (100.0%) | 21 (100.0%) |

Continuous variables are presented as median (interquartile range - IQR). Categorical variables are presented as numbers (n) with percentages (%). Continuous variables for three groups were tested using the Kruskal-Wallis test, followed by Dunn's post-hoc test with Bonferroni correction. Categorical variables were tested using the Chi-square test with Yates' continuity correction. C19 - COVID-19, ARD - Anesthesiology and resuscitation department, ICU - Intensive care unit. (\* $P < 0.05$ for Post-mild C19 vs Post-severe C19, \*\* $P < 0.01$ for Post-mild C19 vs Post-severe C19, \*\*\* $P < 0.001$ for Post-mild C19 vs Post-severe C19, \$\$\$ $P < 0.001$ for Post-mild C19 vs Post-critical C19). C19 – COVID-19.

**Table 2:** Demographic characteristics of sepsis patients.

| RECRUITED SEPTIC PATIENTS |  | n=64 |
| --- | --- | --- |
| Age | Median (IQR) | 71.0 (60.5; 77.0) |
| Sex |  |  |
|  | Male (n (%)) | 44 (68.8%) |
|  | Female (n (%)) | 20 (31.2%) |
| BMI | Median (IQR) | 28.6 (24.7; 32.40) |
| Charlson comorbidity index | Median (IQR) | 4.0 (3.0; 6.0) |
| Number of comorbidities |  |  |
|  | 0 (n (%)) | 12 (18.8%) |
|  | 1 (n (%)) | 19 (29.7%) |
|  | 2 (n (%)) | 13 (20.3%) |
|  | 3 (n (%)) | 10 (15.6%) |
|  | 4 (n (%)) | 5 (7.8%) |
|  | 5 (n (%)) | 3 (4.7%) |
|  | 6 (n (%)) | 2 (3.1%) |
| Smoking |  |  |
|  | Current smoker (n (%)) | 22 (34.4%) |
|  | Ex-smoker (n (%)) | 12 (18.8%) |
|  | Non-smoker (n (%)) | 23 (35.9%) |
|  | NA (n (%)) | 7 (10.9%) |
| SOFA | Median (IQR) | 10.0 (8.0; 12.0) |
| CRP | Median (IQR) | 255.6 (161.8; 335.5) |
| PCT (n=21) | Median (IQR) | 6.5 (1.6; 79.7) |
| Leu |  |  |
|  | Median (IQR) | 14.9 (10.3; 21.7) |
| Catecholamine dose (standard dilution) |  |  |
|  | Median (IQR) | 5.5 (2.0; 12.0) |
| Catecholamine dose (recalculated) |  |  |
|  | Median (IQR) | 0.104 (0.037; 0.269) |
| Lactate |  |  |
|  | Median (IQR) | 1.7 (1.3; 2.9) |
| Bilirubin (n=53) |  |  |
|  | Median (IQR) | 14.3 (8.6; 22.9) |
| Troponin |  |  |
|  | Median (IQR) | 186.0 (113.0; 298.0) |
| Kreatinin |  |  |
|  | Median (IQR) | 133.0 (92.0; 219.8) |
| FiO<sub>2</sub> |  |  |
|  | Median (IQR) | 40.0 (30.0; 55.0) |
| paO<sub>2</sub> |  |  |
|  | Median (IQR) | 12.0 (9.9; 13.3) |
| Horowitz |  |  |
|  | Median (IQR) | 163.5 (90.4; 263.2) |
| Source of sepsis |  |  |
| Abdominal infection (n (%)) |  | 12 (18.8%) |
| Cholangitis (n (%)) |  | 2 (3.1%) |
| Soft tissue infection (n (%)) |  | 5 (7.8%) |
| Neuroinfect (n (%)) |  | 4 (6.3%) |
| Pneumonia (n (%)) |  | 27 (42.2%) |
| Pneumonia + COVID-19 (n (%)) |  | 3 (4.7%) |
| Urosepsis (n (%)) |  | 9 (14.1%) |
| Other (n (%)) |  | 2 (3.1%) |
| Anti-BCG IgG |  |  |
| Negative (n (%)) |  | 40 (62.5%) |
| Positive (n (%)) |  | 24 (37.5%) |
|  | Median (IQR) | 0.8 (0.6; 1.2) |
| Anti-CMV IgG |  |  |
| Negative (n (%)) |  | 9 (14.1%) |
| Positive (n (%)) |  | 55 (85.9%) |
|  | Median (IQR) | 110.3 (32.1; 171.9) |
| Anti-*T. gondii* IgG |  |  |
| Negative (n (%)) |  | 36 (56.2%) |
| Positive (n (%)) |  | 28 (43.8%) |
|  | Median (IQR) | 0.32 (0.16; 2.58) |

Continuous variables are presented as median (interquartile range - IQR). Categorical variables are presented as numbers (n) with percentages (%). BMI – body mass index, SOFA - Sequential Organ Failure Assessment, CRP – C reactive protein, PCT – procalcitonin, FiO<sub>2</sub> - The fraction of inspired oxygen, PaO<sub>2</sub> - the partial pressure of oxygen in the arterial blood, CMV – cytomegalovirus, BCG – Bacillus Calmet-Guerin, *T. gondii* – Toxoplasma gondii.

### Post-COVID-19 patient cohort

Adult patients hospitalised at St. Anne’s University Hospital in Brno, Czech Republic, with severe to critical COVID-19 between October 2021 and September 2024, as well as non-hospitalized volunteers with moderate COVID-19, were recruited 3–17 months (median=6) after infection. Patients undergoing ongoing chronic immuno-suppression therapy or oncological disorders were excluded from the study. SARS-CoV-2 variants were not identified. Written informed consent was obtained from all participants, and all procedures were approved by the institutional ethics committee of St. Anne’s University Hospital Brno (10G/2021).

### Sepsis patient cohort

Adult patients hospitalised at Intensive Care Unit (ICU) of St. Anne’s University Hospital in Brno, Czech Republic, with early septic shock were enrolled. Patients with chronic immuno-suppression therapy or active oncological disease were excluded. Written informed consent was obtained from all participants, and all procedures were approved by the institutional ethics committee of St. Anne’s University Hospital Brno (10G/2021).

### BCG-vaccinated and non-vaccinated volunteer cohort

Adult volunteers vaccinated or not with BCG during childhood were recruited between August 2022 and July 2023 (BCG vaccinated: n=8, median age [min-max] = 34 [28–41], sex [men/women] = 4/4; BCG non-vaccinated: n=8, median age [min-max] = 27 [23–34], sex [men/women] = 4/4). Patients with chronic immuno-suppression therapy or active oncological disease were excluded. Written informed consent was obtained from all participants, and all procedures were approved by the institutional ethics committee of St. Anne’s University Hospital Brno (10G/2021).

### Control cohort

An age- and comorbidity-matched control cohort, consisting of individuals hospitalised at St. Anne’s University Hospital in Brno one day before elective orthopaedic surgery (n=10; median age [min-max] = 75 [64–83]; sex [men/women] = 5/5; median BMI [min-max] = 29.2 [23.5– 35.9]; median Charlson comorbidity index [min-max] = 3.5 [2–8]) was also included. As a positive control for activation-induced markers, we used stabilised blood samples from severe COVID-19 patients in the acute phase (within 15 days of hospitalisation; n=8; sex [men/women] = 4/4; median age [min-max] = 59 [41–85]; median BMI [min-max] = 30 [23–39]; median Charlson comorbidity index [min-max] = 2 [0–6]), obtained from our previous study.[37] Written informed consents were obtained from all participants, and all procedures and protocols were approved by the institutional ethics committee of St. Anne’s University Hospital Brno (control cohort: 11G/2021; acute COVID-19: 6G/2022).

### Blood sample processing and plasma preparation

Blood samples were processed within 2 hours of collection. Stabilised blood samples were prepared by incubating 0.2 mL of whole blood with an equal volume of Whole Blood Cell Stabilizer (Cytodelics AB, Sweden) at room temperature for 15 minutes, then stored at −80°C until processing.[37] Plasma was separated by centrifuging whole blood at 2500 g for 15 minutes at 4°C and immediately frozen at −80°C.

### Detection of anti-CMV, anti-BCG, anti-T. gondii

Anti-CMV, anti-BCG, and anti-*T. gondii* IgG levels in plasma were measured using commercial ELISA kits (Cytomegalovirus IgG ELISA Kit [Creative Diagnostics, DEIA326], Human Anti-Tuberculosis [BCG] IgG and Ig ELISA kit [Alpha Diagnostic Intl. Inc.], and EIA Toxoplasma [TestLine Clinical Diagnostics]) according to the manufacturer’s instructions.

### Immunophenotyping of stabilized whole blood

Stabilised blood samples were processed as described above. Fc receptor (FcR) binding sites were blocked with FcR blocking solution (Miltenyi Biotec, Auburn, CA, USA) for 15 min at 4°C. Samples were stained with antibody cocktails (Supplementary Table 1) for 30 min at 4°C. Sample acquisition was performed on the BD FACSymphony™ A1 (BD Biosciences). To account for batch effects, a common sample was included in each batch, and only markers without batch effects were used for downstream analysis. Data were analysed using FlowJo® v10.10 (FlowJo, LLC, Ashland, OR, USA). Gating strategies are presented in Supplementary Fig. 1.

### Statistical analysis

Prism® (GraphPad Software LLC Ltd, La Jolla, CA, USA) software and R v4.2.3. (R Core Team 2021, R Foundation for Statistical Computing, Vienna, Austria) were used for statistical analyses. Data were tested for normality using Shapiro-Wilk test and graphically by Q-Q plots and histograms. Continuous variables were presented as median (interquartile range). Categorical variables were presented as numbers with percentages. Continuous variables for 3 or more groups were tested using the Kruskal-Wallis test, followed by Dunn’s post-hoc test with Bonferroni correction. For comparisons done between two groups, the Wilcoxon test was used. Categorical variables were tested using the Chi-square test with Yates’ continuity correction. Correlation was calculated using Spearman’s correlation coefficient. For comparing dependent variables, the Friedman test was applied, followed by pairwise Wilcoxon post-hoc tests with Bonferroni correction. Any deviation from the abovementioned statistical tests is described in the figure legend or the appropriate section of the results. The level of statistical significance was determined as follows: * (*P* <0.05), ** (*P* <0.01), *** (*P* <0.001), and **** (*P* <0.0001).

### Monocyte isolation from fresh blood

Peripheral blood from post-COVID-19 patients was collected in Li-heparin tubes. Monocytes were isolated using the RosetteSep Human Monocyte Enrichment Cocktail (STEMCELL Technologies, Vancouver, Canada) and gradient centrifugation with Lymphoprep (density 1.077 g/mL; STEMCELL Technologies) according to the manufacturer’s recommendations. Isolated monocytes were cryopreserved until use.

### Monocyte/HSPCs enrichment from PBMCs

PBMCs were isolated from blood of BCG-vaccinated and non-vaccinated volunteers using gradient centrifugation with Lymphoprep (density 1.077 g/mL; STEMCELL Technologies). Monocytes were enriched with CD14+ microbeads (MACS, Miltenyi), and CD34+ hematopoietic stem and progenitor cells (HSPCs) were enriched from the negative fractions using CD34 microbeads kit ultra-pure (MACS, Miltenyi). Isolated cells were used for stimulation assays and methylcellulose colony-forming unit (CFU) assays.

### Monocyte stimulation

Monocytes were seeded at 1 × 10^5^ cells per well in RPMI with 10% FBS and incubated overnight at 37°C with 5% CO₂. Cells were then stimulated with LPS (1 µg/mL LPS from *E. coli* O55, InvivoGen), SARS-CoV-2 (omicron variant B.1.1. 529, isolate: 17577/21, MOI 0.1), opsonised SARS-CoV-2 (VNT serum, titer 1:320, 1 hr at 37 °C), inactivated SARS-CoV-2 using 0.1% β-propiolactone (overnight incubation at 2-8 °C, MOI 0.1), or zymosan (zymosan from *Saccharomyces cerevisiae*, InvivoGen, 5 µg/mL) for 24 h. Harvested cells were lysed in TriReagent (Molecular Research Center) for RNA isolation.

### RNA isolation and gene expression analysis

RNA was extracted from monocytes using TriReagent (Molecular Research Center) and purified with RNeasy spin columns (Qiagen, Hilden, Germany). RNA concentration and quality were assessed using a NanoDrop spectrophotometer (Agilent, Santa Clara, CA, USA). RNA was transcribed into cDNA using the High-Capacity cDNA Reverse Transcription Kit (Thermo Fisher Scientific). qPCR was performed with TaqMan Gene Expression Assays (Thermo Fisher Scientific) on a LightCycler II (Roche, Basel, Switzerland). Relative gene expression was calculated as 2−ΔCt, normalising to GAPDH. TaqMan probes used included TNF-ɑ (Hs00174128_m1), IL-1β (Hs01555410_m1), IL-6 (Hs00174131_m1), and GAPDH (Hs02758991_g1).

### Methylcellulose CFU assay

CD34+ cells (3 × 10^2^ cells/mL) were seeded in methylcellulose media (MethoCult™ GF+ H4435, STEMCELL Technologies) and incubated at 37°C with 5% CO₂ for 16 days. Colonies were counted, harvested, washed, and stained with antibodies targeting CD34 (APC-eF780, clone 4H11, eBioscience), CD11b (PE-Cy7, clone ICRF44, Biolegend), CD13 (APC, clone WM-15, eBioscience), CD15 (eF450, clone HI98, eBioscience) CD235ɑ (PE, HIR2 [GA-R2], Invitrogen), CD14 (BV510, clone M5E2, Sony), CD45 (PerCP-eF710, clone HI30, Invitrogen), and a LIVE/DEAD fixable dead stain kit (Green, Invitrogen). Data was acquired on FACS Canto II (BD Biosciences), and analysed with FlowJo® v10.10. Gating strategies are presented in Supplementary Fig. 1.

## RESULTS

### Immune dysregulation signs are present in myeloid and lymphoid immune cells of post-COVID-19 patients

To investigate the effect of COVID-19 on the immune system, we performed whole blood immunophenotyping and monitored the changes in frequencies and functionality of immune cells according to disease severity (Supplementary figure 2). First, we focused on changes in the myeloid cell compartment in post-COVID-19-patients (mild n=41, severe n=27, critical n=21), acute COVID-19 patients (n=8) and age-matched controls (n=10). FACS analysis of whole blood samples showed dysregulation in the frequency of total neutrophils (CD45^+^CD3^-^ CD56^-^CD66b^+^) after COVID-19, particularly in post-mild COVID-19 patients compared the acute COVID-19 patients and age-matched controls (Fig. 1A). In contrast, the frequency of total monocytes (CD45^+^CD3^-^CD56^-^CD66b^-^CD14^+/dim^CD16^+/-^) remained unaffected (Fig. 1A). Interestingly, at the monocyte subset level, we observed a reduction in classical monocytes (CD14^+^CD16^-^) and an increase in non-classical (CD14^low^CD16^+^) monocytes in post-severe COVID-19 patients compared to post-mild COVID-19 group (Fig. 1B), raising the possibility that COVID-19 severity may be linked to long-term alterations in monocyte populations.

**Figure 1:**
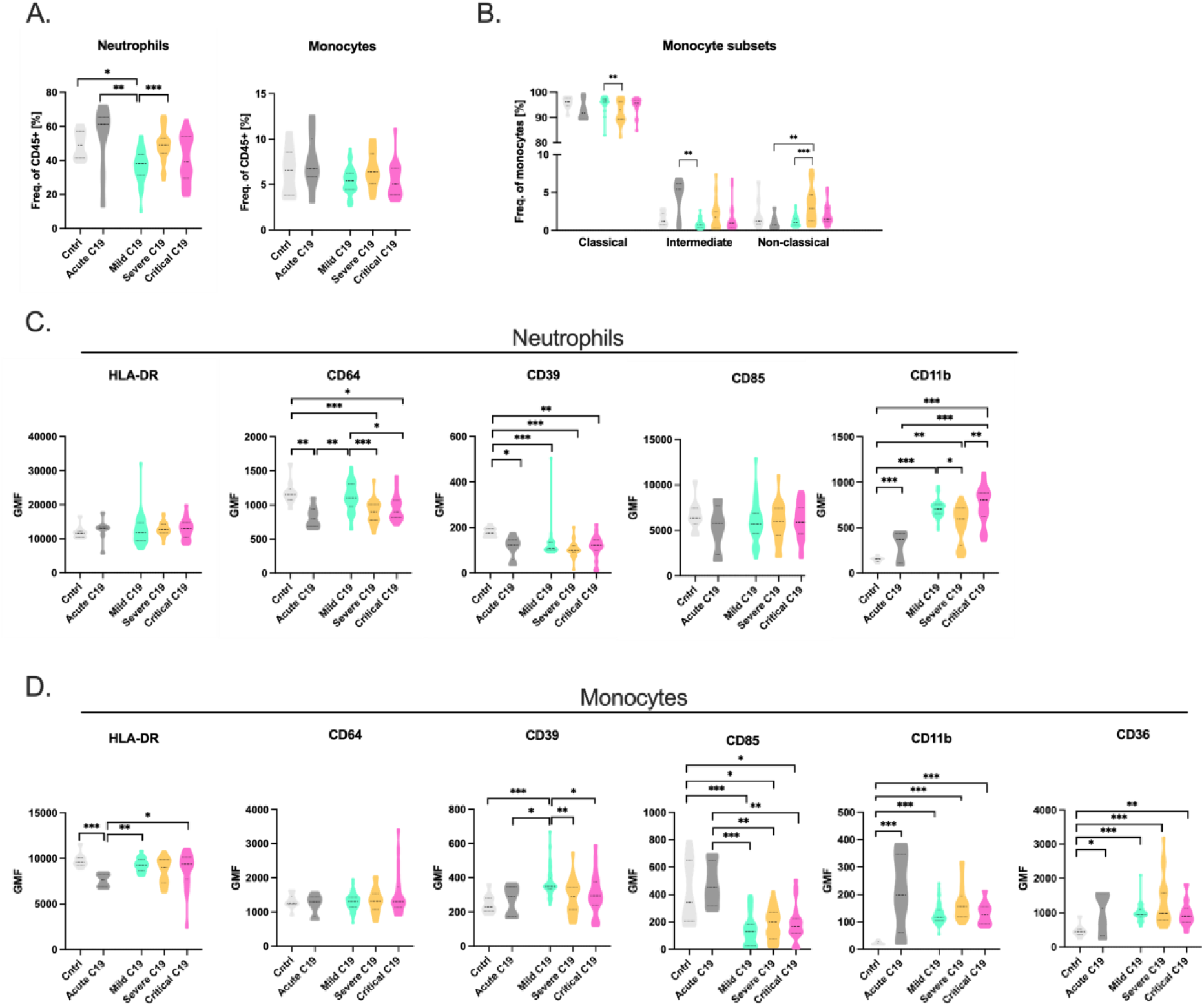
Long-term impact of COVID-19 and its severity on myeloid immune cells. A.) Changes in frequencies of monocytes and neutrophils observed in patients who recovered from mild (n=41), severe (n=27), critical (n=21) COVID-19. An age- and comorbidity-matched control cohort (Cntrl, n=10) and acute COVID-19 patients (Acute C19, n=8) were also included. B.) Frequency of monocyte subsets: classical (CD14^+^CD16^-^), intermediate (CD14^+^CD16^+^) and non-classical (CD14^low^CD16^+^) monocytes found in post-mild, -severe, -critical COVID-19 patients. C.) Expression levels of functionality-related markers in neutrophils. D. Expression levels of functionality-related markers in monocytes. Data were tested using Kruskal-Wallis test followed by post-hoc Dunn’s test with Bonferroni correction. Statistically significant differences are indicated as follows: *P < 0.05, **P < 0.01, ***P < 0.001.

To determine how COVID-19 severity affects the activation and functional status of monocytes and neutrophils, we performed a detailed characterization based on expression of relevant immune-cell activation and functional markers. For function, we evaluated the expression of CD36, CD11b, CD85d, CD39, CD64 and HLA-DR. In neutrophils (Fig. 1C), we detected significantly increased CD11b expression in all post-COVID-19 patients compared to controls and acute COVID-19 patients. By contrast, CD39 was significantly decreased when comparing post-COVID-19 patients with controls. Finally, CD64 expression was significantly decreased in post-severe and post-critical COVID-19 patients compared to post-mild COVID-19 patients and controls. In monocytes (Fig. 1D), we observed significantly increased CD36 and CD11b expression and decreased CD85d expression in all post-COVID-19 patients when compared to controls. Similarly, CD39 was significantly increased on monocytes from post-mild COVID-19 patients compared to controls. CD64 and HLA-DR remained unchanged when compared to controls. These data suggest that dysregulated myeloid cells persist in post-COVID-19 patients regardless of COVID-19 severity.

Having detected persistent alterations of myeloid cells in post-COVID-19 patients, we next aimed to deeply characterize the impact of COVID-19 severity on lymphoid cell frequencies and function by FACS analysis (Supplementary figure 2). We showed that the frequencies of total NK cells (CD45^+^CD3^-^CD56^+^) and T cells (CD45^+^CD3^+^) were dysregulated during the acute state of COVID-19 (n=8) compared to post-COVID-19 patients (mild n=41, severe n= 35, critical n=21) and age-matched controls (n=10) (Fig. 2A-B). In agreement with our previous observation [37], the frequencies of NK cells and T-cells normalised in majority of convalescent patients several months after COVID-19 (Fig. 2A). However, we noticed persistently reduced NK cell frequencies in post-critical COVID-19 patients in comparison to controls. In addition, we evaluated the frequencies of specific immune cell subset (Fig. 2C-D). We observed a significantly increased frequency of CD56^dim^CD16^+^ NK cells in all post-COVID-19 groups in comparison to the acute COVID-19 and the controls. In T cells, we showed persistently reduced CD4^+^, Tregs (CD4^+^CD127^-^CD25^+^) and increased CD8^+^ in post-critical COVID-19 compared to post-mild COVID-19. Looking specifically at CD8^+^ T cells, we also reported an increased frequency of CD57^+^CD45RA^+^ CD8^+^ T cells in post-critical COVID-19 in comparison to post-mild/severe COVID-19 patients.

**Figure 2:**
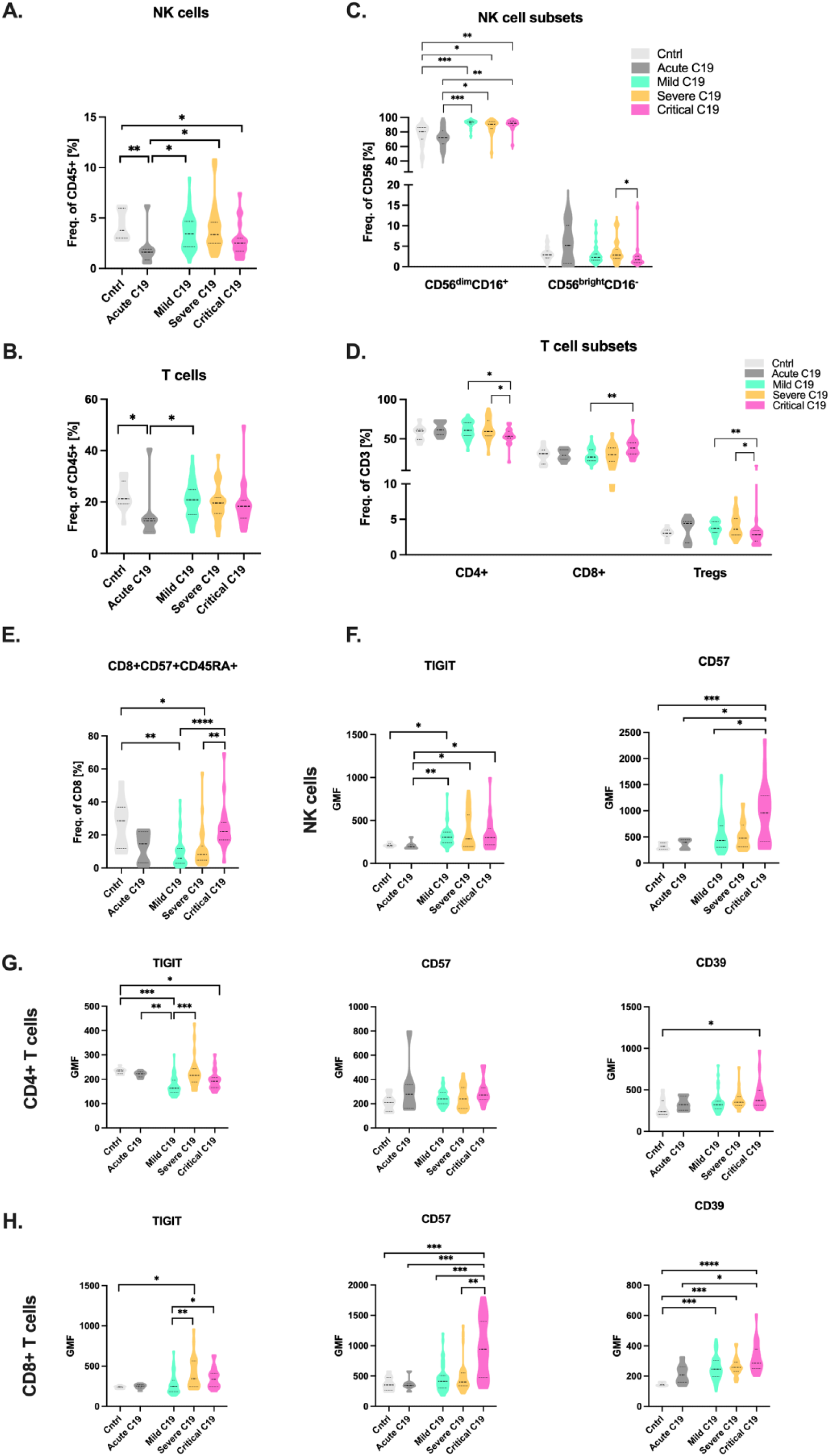
Long-term impact of COVID-19 and its severity on lymphoid cells. Changes in frequencies of A.) NK cells and B.) T cells observed in patients who recovered from mild (n=41), severe (n=27), critical (n=21) COVID-19. An age- and comorbidity-matched control cohort (Cntrl, n=10) and acute COVID-19 patients (Acute C19, n=8) were also included. COVID-19. Frequency of main C.) NK cell and D.) T cell subsets, and CD8^+^CD57^+^CD45RA^+^ T cells (E.) in post-mild/severe/critical COVID-19 patients. Expression levels of relevant functionality-related markers in F.) NK cells, G.) CD4+ Tcells and H.) CD8+ T cells. Data were tested using Kruskal-Wallis test followed by post-hoc Dunn’s test with Bonferroni correction. Statistically significant differences are indicated as follows: *P < 0.05, **P < 0.01, ***P < 0.001, ****P < 0.0001. Abbreviation: Cntrl – Control group; C19 - COVID-19.

Taken together these results showed alterations in frequencies of NK and T cells in convalescent patients. In particular, we observed persistently lower total NK cell counts accompanied by the increase in CD56dimCD16+ NK cell subset characterized by potent cytotoxic functions, suggesting a compensatory mechanism to restore NK cell function. We also demonstrated COVID-19 severity-dependent impact on frequencies of CD4, Tregs and CD8 T cells. However, further investigation will be needed to precisely determine the implications of these results.

To further evaluate activation and exhaustion status of lymphoid cells, we focused on expression of CD57, TIGIT, and CD39. In NK cells (Fig. 2F), CD57 expression was significantly increased in post-critical COVID-19 patients compared to post-mild-COVID-19, acute-COVID-19, and control groups. Meanwhile, TIGIT expression was significantly increased in all post-COVID-19 groups compared to acute COVID-19. However, we found persistently increased TIGIT expression only in post-mild COVID-19 patients compared to controls. Overall, the higher expression of CD57 and TIGIT on NK cells in all convalescent COVID-19 patients might indicate NK cells exhaustion and inhibition of their functions.

In CD4^+^ T-cells (Fig. 2G), we observed significantly lower TIGIT expression in post-mild and post-critical COVID-19 patients compared to acute COVID-19 patients and controls, while CD39 expression was significantly increased in post-critical COVID-19 patients compared to controls. CD57 expression was unaffected. Finally, we analysed CD8^+^ T cells (Fig. 2H), where we detected significantly increased CD57 expression in post-critical COVID-19 patients compared to all other groups, as well as TIGIT in post-severe and post-critical COVID-19 patients compared to post-mild COVID-19 patients. Meanwhile, CD39 was significantly increased in all post-COVID-19 patients compared to controls. The altered expression of these markers in T cells, particularly in CD8+, implies an immunosuppressive state of these populations.

Overall, NK and T cells seem to be influenced by previous COVID-19 infection, most notably the CD8 compartment where perturbed TIGIT, CD39 and CD57 expression suggests dysfunctional immune status and a possible link to exhaustion and immunosuppression.

### Association of plasma anti-BCG, *T. gondii,* and CMV IgG levels with COVID-19 severity and sepsis survivorship

Based on extensive studies, BCG vaccination is suggested to provide non-specific protection against unrelated pathogens. Here, we aimed to determine whether BCG seropositivity as well as latent infections (CMV, *T. gondii*) are associated with COVID-19 severity. Since COVID-19 shares many similarities with sepsis, as evidenced by the fact that critically ill COVID-19 patients meet the Sepsis-3 criteria[38] and present infection-associated organ dysfunction,[39] we also aimed to investigate and compare the seroprevalence of BCG, CMV, and *T. gondii* in a cohort of sepsis patients. To do so, we recruited 97 patients who had recovered from mild to critical COVID-19 (mild n=41, severe n= 35, critical n=21) (Tab. 1) and 64 patients with sepsis (Tab. 2). From these cohorts, we first aimed to investigate the relationship between persistent plasma anti-BCG IgG levels and latent infections (*T. gondii*, CMV) with COVID-19 severity. We found significant differences in the BCG seropositivity between the COVID-19 patients’ groups (Fig. 3A). Specifically, more patients with undetectable anti-BCG in plasma were found in the severe and critical COVID-19 groups than in the mild COVID-19 group (60.0%, 57.1%, and 34.1% respectively). We also observed a significantly higher frequency of patients with latent *T. gondii* infection in the severe (28.6%) and critical (42.9%) COVID-19 patient groups versus the mild COVID-19 patient group (9.8 %). We saw a similar pattern for those with a latent CMV infection (68.6% and 95.2% versus 58.5%) (Fig. 3A). In the sepsis cohort, the frequency of patients seropositive for BCG, CMV, and *T. gondii* mirrored the rates observed in critically ill COVID-19 patients. In addition, we found that sepsis patients positive for anti-BCG IgG in the plasma (IgG level > 3U/ml) had higher 28-days survival than patients negative for BCG antibodies (*P* = 0,059; Fig. 3C). By contrast, we saw no significant differences in survival outcomes among the groups positive or negative for plasma *T. gondii* and CMV antibodies (Fig. 3C). Of note, a further analysis identified a significant positive correlation between CMV and *T. gondii* IgG levels and age (Fig. 3B) but not between BCG positivity and age (Fig. 3B). Thus, it is plausible that BCG seropositivity may be linked to milder COVID-19 and improved survival outcomes sepsis.

**Figure 3:**
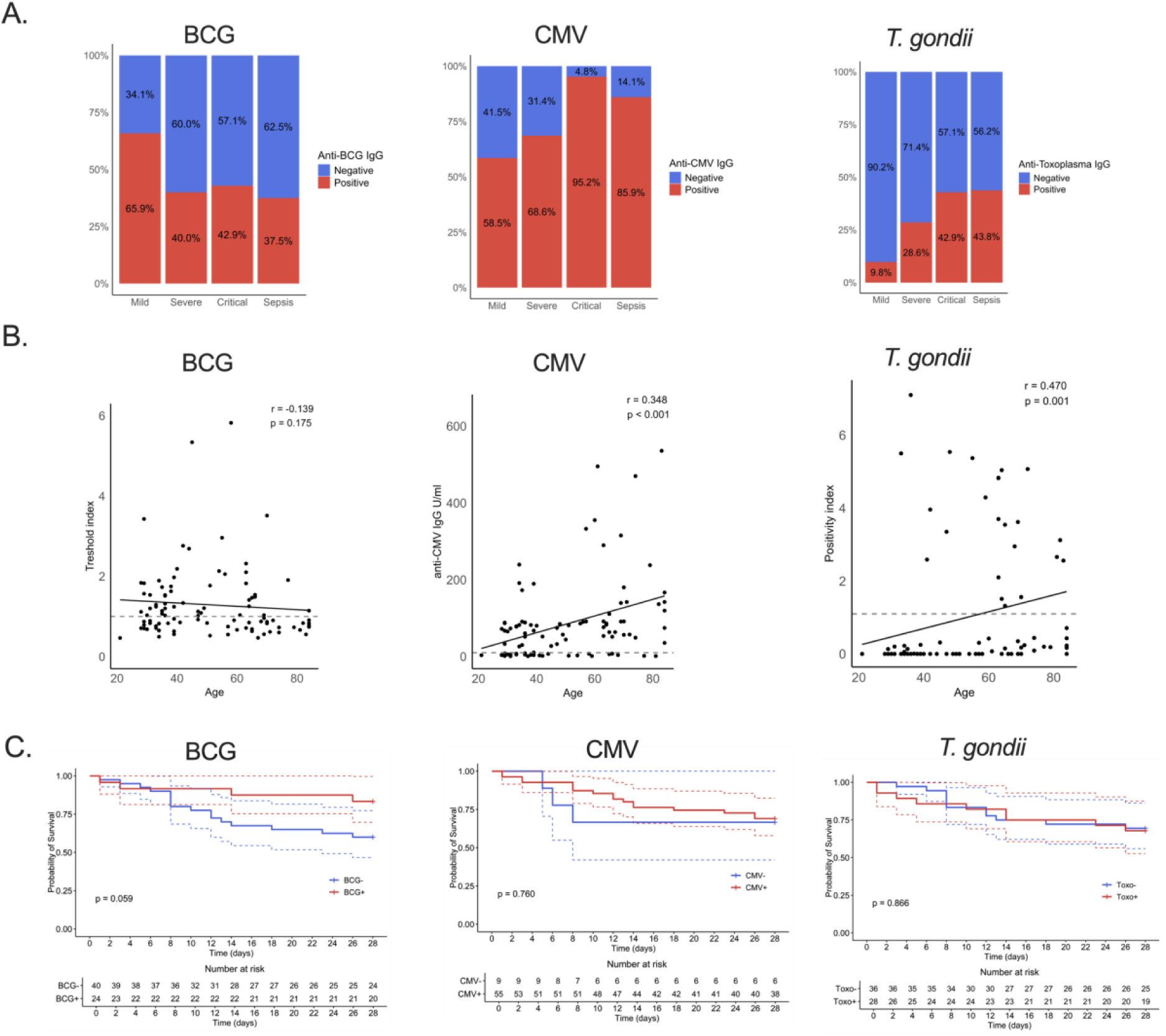
Persistent antibody levels long after BCG vaccination are associated with milder COVID-19 progression and better survival prognosis in sepsis. A.) Seroprevalence of anti-BCG/CMV/*T. gondii* IgG in patients who recovered from mild/severe/critical COVID-19 and in sepsis cohort. B.) Correlation between anti-BCG/CMV/T*. gondii* IgG levels and age of post-COVID-19 patients (Spearman’s correlation). C.) Kaplan-Meier survival curves (solid lines) and 95% confidence intervals for the group of sepsis patients with detectable levels of anti-BCG/CMV/*T. gondii* IgG (red) and the group with undetectable levels (blue). Log-rank test was used to test statistical significance. Abbreviation: BCG – Bacillus Calmet Guérin; CMV -cytomegalovirus; Cntrl – Control group; *T. gondii* – *Toxoplasma gondii*; C19 - COVID-19.

### Monocyte response to SARS-CoV remains unaffected by previous COVID-19 or BCG

Given the observed altered expression of activation and functional markers by monocytes and a potential association of BCG seroprevalence with better disease outcomes, we next wanted to understand whether the monocyte response to microbial challenge is affected. To do so, we isolated monocytes from post-COVID-19 patients (n=24) 4-10 months (median = 8) after COVID-19, and stimulated them with LPS or SARS-CoV-2. Monocytes significantly upregulated IL-1β and IL-6 gene expression after 24 h of stimulation with either stimulus (Fig. 4A) suggesting that the ability of monocytes to respond to microbial stimuli remains intact post COVID-19. The magnitude of *ex vivo* response of patients’ monocyte to stimulation was not affected by COVID-19 severity (Supplementary Fig. 3A).

**Figure 4:**
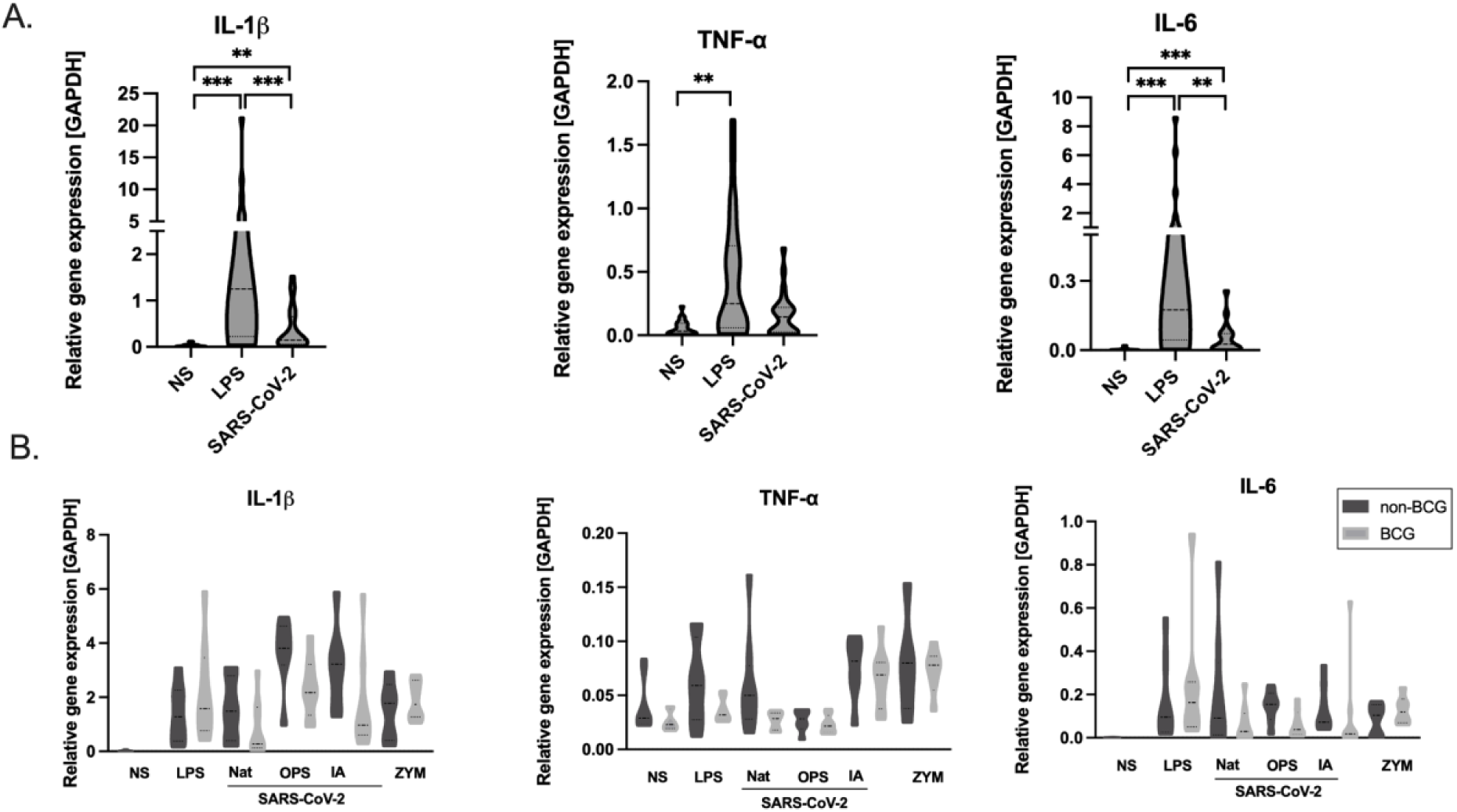
Long-term impact of COVID-19 and BCG vaccination on monocyte response to microbial triggers measured at mRNA level. A.) Inflammatory genes expression measured by qPCR in post-COVID-19 patients triggered with LPS and SARS-CoV-2 B.) Impact of BCG vaccination in childhood on monocyte response to LPS, native-(NAT), opsonized-(OPS), inactivated-(IA) SARS-CoV-2, and zymosan (ZYM). Statistics: Kruskal Wallis + Dunn’s multiple comparison test. Statistically significant differences are indicated as follows: *P < 0.05, **P < 0.01, ***P < 0.001. *non-BCG - individuals without previous BCG vaccination, BCG – individuals vaccinated with BCG in childhood*.

To investigate the possible link between BCG-induced unspecific protection against SARS-CoV-2 several early epidemiological studies found that inhabitants of countries lacking a BCG vaccination policy were more prone to SARS-CoV-2 infection and succumbing to COVID-19. Thus, we aimed to determine association between previous BCG vaccination given in childhood and the monocyte response to microbial challenge. Because monocytes are short-lived immune cells, the long-term effects of TI arise from reprogramming of progenitor cells [14]. Therefore we simultaneously isolated monocytes and CD34^+^ hematopoietic stem and progenitor cells (HSPCs) from peripheral blood mononuclear cells (PBMCs) of BCG-vaccinated (n=8) and non-vaccinated (n=8) individuals. We stimulated isolated monocytes with either native, opsonized, or inactivated SARS-CoV-2, LPS, or zymosan and measured IL-6, IL-1β and TNF-ɑ expression levels. We found no differences in the monocyte response to any stimuli when comparing BCG-vaccinated and non-vaccinated groups (Fig. 4B). However, when we evaluated the monocyte response regardless of the BCG vaccination, we found that TNF-ɑ was significantly upregulated by monocytes exposed to inactivated SARS-CoV-2 but not opsonized SARS-CoV-2 (Supplementary Fig 3B). This finding implies that mechanism of virus recognition could be important in shaping the magnitude of the immune response.

Finally, we were interested in determining the differentiation capacity of isolated HSPCs. We saw no significant differences in terms of numbers of the colonies formed (Supplementary Figure 4A-4B) or their phenotypes (Supplementary Figure 4C) when comparing the HSCPs from the vaccinated versus unvaccinated individuals. As such, we can conclude that the hypothesized TI effect of BCG vaccination administered in childhood is not preserved in adulthood.

Overall, our results revealed long-term alterations of myeloid and lymphoid immune cells in post-COVID-19 patients underscoring a profound long-term impact of COVID-19 on immune system. Furthermore, we observed lower BCG seropositivity together with higher rate of latent infections (CMV, BCG) in severe/critical-post-COVID-19 patients as well as sepsis patients. However, our data showed no relationship between previous BCG vaccination and protection against SARS-CoV-2 infection.

## DISCUSSION

Currently, the understanding of how COVID-19 severity impacts long term immunity remains limited. Here, we provide a comprehensive characterization of immune cells in the patients who recovered from mild-, severe- and critical-COVID-19. We observed that immune dysregulation persisted months after COVID-19. While our analysis showed only minor changes in frequencies of innate cells among the post-COVID-19 cohorts, the detailed analysis of activation markers revealed persistent differences. Specifically, profound changes in neutrophil population amongst severe COVID-19 patients were associated with increased CD11b and decreased CD39 and CD64 expression. These changes in CD11b indicate dysregulation in phagocytic capacity as well as in adhesion, and migration [40], while decreased CD39 indicates alleviation of anti-inflammatory potential [41]. We hypothesize that reduced CD64 expression on neutrophils could be associated with receptor shedding, since this receptor was reported as highly upregulated during acute COVID-19 [42].

Monocytes from all COVID-19 patients exhibited dysregulated expression of several markers in comparison to controls. Specifically, increased expression of CD11b suggests an impact on phagocytic function, adhesin and migration and elevated CD36 expression points toward impaired lipid metabolism [42] and senescence [43], while decreased CD85d might indicate persistent activation status [44]. Notably, rescued HLA-DR expression in convalescent patients indicates functional antigen presentation capacity post-COVID-19 regardless of disease severity.

We have also analysed the phenotype of adaptive immune cells showing long-term alterations. T cells normalized in majority of convalescent patients within several months, which aligns with our previous findings [37]. However, the frequency of NK cells remained lower in patients, who recovered from critical COVID-19. Similarly, reduction in NK cell frequencies during acute COVID-19 was followed by slow recovery in post-COVID-19 patients [45]. In addition, we showed persistently higher frequency of CD56^dim^CD16^+^ NK cells across all post-COVID-19 patients which is in agreement with a recent study [45]. Regarding T cell subsets, we reported lower CD4^+^ and Tregs shifting towards increased CD8^+^ T cell frequencies in post-critical compared to post-mild COVID-19 patients aligning with studies showing vital role of CD8^+^ T cells in COVID-19 resolution [46, 47].

Furthermore, we observed changes in the expression of activation and exhaustion markers in lymphoid cells. Elevated expression of CD57, TIGIT and CD39 on CD8^+^ T cells suggests terminal differentiation and exhaustion, indicative of potential immune dysfunction [48-50]. Notably, CD57^+^ expression on CD8^+^ T cells was exclusively elevated in post-critical COVID-19 patients in comparison to post-mild and severe patients indicating a significant impact of disease severity on CD8^+^ T cells and their function. In addition, these findings suggest a possible link with enhanced immune aging in survivors of critical COVID-19 [42].

While the role of BCG in TI is well established [37], the possible contribution of CMV and *T. gondii* infections to TI development remains elusive. We observed differences in seropositivity for CMV, *T. gondii* and BCG antibodies across all studied cohorts. Interestingly, patients who suffered from severe or critical COVID-19 were most frequently positive for CMV and *T. gondii* yet they exhibited the lowest BCG seropositivity. We observed similar trend in CMV, *T. gondii* and BCG seropositivity in sepsis patients. Further analysis revealed a positive correlation between CMV or *T. gondii* positivity and age, confirming previous reports [51-54]. However, we could not distinguish between the impact caused by disease severity and age. With the overall seroprevalence of *T. gondii* in the Czech Republic reported at ∼23-32 % [53, 55], we speculate that latent *T. gondii* infection may similarly influence COVID-19 severity, as previously noted by Flegr et al. [56] though indirect effects cannot be excluded [57]. CMV and *T. gondii* seroprevalence varies by country and age [51-54], reaching 50-90% [51] and 10-80% globally [58, 59], respectively. Thus, throughout the lifetime, CMV affects most individuals, causing lifelong latent infection with occasional episodes of reinfections, and *T. gondii* follows a similar aetiology. In contrast, we showed that anti-BCG IgG levels did not correlate with age, indicating a potential age-independent protective effect. Rather, our data support that BCG seropositivity is associated with improved survival in sepsis patients, consistent with findings that BCG vaccination may protect against severe neonatal sepsis [57, 60, 61]. However, similar studies in adults are lacking, and larger clinical trials are needed to confirm these observations and inform preventive strategies. Both infections (CMV, *T. gondii*) have been studied in the context of sepsis severity [7, 9, 59] or TI [62] and also for their potential risk of reactivation and reinfection of vulnerable immunosuppressed sepsis patients [63]. Moreover, CMV has a profound impact on immune system composition and function, including the accumulation of CMV-specific memory T cells (memory inflation), particularly terminally differentiated T cells associated with immunosenescence [51].

To assess the impact of TI induced by SARS-CoV-2 and BCG vaccination, we exposed monocytes from post-COVID-19 patients (mild n=8, severe n=5, critical n=11, median time after COVID-19 = 8 months), BCG-vaccinated (n=8) and unvaccinated volunteers (n=8) to LPS, zymosan, and various forms of SARS-CoV-2. No significant differences in IL-1β, TNF-α, or IL-6 production were observed in response to these stimuli, suggesting that neither COVID-19 nor previous BCG vaccination have effect on monocyte responsiveness. Our findings are supported by a recent study showing that administration of BCG vaccine at birth does not have a protective effect against COVID-19 [64]. Interestingly, TNF-α levels were elevated in monocytes exposed to inactivated versus opsonized SARS-CoV-2, emphasising the importance of SARS-CoV-2 recognition pathways in monocyte responses. This finding might be particularly relevant for vaccine development [65].

This single-centre observational study has also some limitations. The major limitation of our study is a relatively small sample size of sub-cohorts based on different severity of COVID-19. Furthermore, the patients were enrolled over a relatively long period (from October 2021 to September 2024), during which different variants of SARS-CoV-2 were circulating, which may have had an impact on the heterogeneity of the (sub)cohorts. We also do not have available clinical data assessing post-COVID-19 syndrome. Further studies evaluating the link between our findings showing immune dysregulation and long COVID-19 are needed. Another limitation is that we assessed the effect of BCG administered in childhood on adults. The long-term duration of innate immune memory (trained immunity) is still not fully understood, but it is currently assumed to last at least several months to years. Nevertheless, a clear advantage of our approach is the correlation of the outcomes directly to the BCG antibody plasma titres, as most published studies to date have relied on information about the vaccination itself without data on persisting antibody titres [23-28, 64].

In conclusion, our findings on persistent immunomodulation following COVID-19 reveal significant changes in both myeloid and lymphoid cells, even in patients who experienced mild disease. These alterations may be especially relevant for all COVID-19 survivors, particularly those suffering from long-term post-COVID-19 sequelae. Given the BCG vaccination and both *T. gondii* and CMV infections are highly relevant modulators of the immune system, we studied their association with COVID-19 severity and sepsis survival. While there are vast variations in the uptake of BCG vaccination and the prevalence of *T. gondii* and CMV in the global population, a large proportion of COVID-19 and sepsis patients will exhibit the consequences of TI. Thus, we conclude that BCG seropositivity might be associated with improved COVID-19 and sepsis outcomes. We hope that these data will initiate productive discussions on how such readily obtained diagnostic information could guide patient risk stratification.

## AUTHOR CONTRIBUTIONS

KB, GB, MH, PB conducted the experiments and acquired the data. KB, MH, GB, IP, ES, MHK, JF participated in the analysis of the data. MH, RP, MDH, MV, VŠ contributed to patient recruitment and clinical sample processing. MH, RP, LO contributed to clinical data collection.

KB, JF and MHK supervised the study and wrote the manuscript. KB, MDZ, MHK and JF acquired the funding and design the study. All authors participated in editing and reviewing the manuscript.

## ACKNOWLEDGEMENTS

The authors would like to thank Dr Jessica Tamanini of Insight Editing London for critically reviewing the manuscript before submission, the technician team of the Center for Translational Medicine, International Clinical Research Center, and involved orthopedic ICU and anesthesiology nurses of St. Anne’s University Hospital and the Department of Anesthesiology and Intensive Care, Brno (Czech Republic) for their help with blood sample collection. A special thanks goes to all study participants enrolled at St. Anne’s University Hospital Brno. The authors thank Dr. Ondřej Pelák of BD Biosciences Czechia for providing access to the BD FACSymphony A1 analyzer and for his guidance and expertise with the instrument. Finally, the authors acknowledge the Dr. Petra Wesela from Biostatistics Core Facility of St. Anne’s University Hospital for her support and assistance with biostatistic analysis in this work.

