## Supplementary information for "Long-term immune changes after COVID-19 and the effect of BCG vaccination and latent infections on disease severity"

RUNNING HEADLINE: COVID-19: Immunomodulation & BCG Impact

Kamila Bendíčková[1,2], Ioanna Papatheodorou[1,2,3], Gabriela Blažková[1], Martin Helán[1,4], Michaela Haláková[1], Petr Bednář[5,6,7,12], Erin Spearing[1], Lucie Obermannová[4], Julie Štíchová[8], Monika Dvořáková Heroldová[9], Tomáš Tomáš[10], Roman Panovský[1,11], Vladimír Šrámek[4], Marco De Zuani[1], Marcela Vlková[8], Daniel Růžek[5,7,12], Marcela Hortová-Kohoutková[1,2], Jan Frič[1,2,13]

[1]International Clinical Research Center, St. Anne's University Hospital, Brno, Czech Republic.

[2] International Clinical Research Center, Faculty of Medicine, Masaryk University, Brno, Czech Republic.

[3] Department of Biology, Faculty of Medicine, Masaryk University, Brno, Czech Republic.

[4] Department of Anesthesiology and Intensive Care, St. Anne's University Hospital and Faculty of Medicine, Masaryk University, Brno, Czech Republic.

[5] Veterinary research institute, Brno, Czech Republic.

[6] Department of Medical Biology, Faculty of Science, University of South Bohemia, Ceske Budejovice, Czech Republic.

[7] Department of Experimental Biology, Faculty of Science, Masaryk University, Czech Republic.

[8] Institute of Clinical Immunology and Allergology, St. Anne's University Hospital and Faculty of Medicine, Masaryk University, Brno, Czech Republic.

[9] Department of Microbiology, Anne's University Hospital and Faculty of Medicine, Masaryk University Brno, Czech Republic.

[10] First Department of Orthopaedic Surgery, St. Anne's University Hospital and Faculty of Medicine, Masaryk University Brno, Czech Republic.

[11] 1st Department of Internal Medicine/Cardioangiology, St. Anne's Faculty Hospital, Faculty of Medicine, Masaryk University Brno, Brno, Czech Republic.

[12] Institute of Parasitology, Biology Centre of the Czech Academy of Sciences, Ceske Budejovice, Czech Republic.

[13] Institute of Hematology and Blood Transfusion, Prague, Czech Republic.

**Supplementary table 1**: Clinical characteristics of patients hospitalised with severe and critical COVID-19.

| CLINICAL CHARACTERISTICS OF POST COVID-19 PATIENTS | | | |
|---|---|---|---|
| | **Post-severe C19 (n=35)** | **Post-critical C19 (n=21** | ***P*-value** |
| X-ray | | | |
| 0 (n (%)) | 18 (51.4%) | 0 (0.0%) | |
| 1 (n (%)) | 8 (22.9%) | 3 (14.3%) | |
| 2 (n (%)) | 5 (14.3%) | 1 (4.8%) | < 0.001 |
| 3 (n (%)) | 2 (5.7%) | 7 (33.3%)t | |
| 4 (n (%)) | 2 (5.7%) | 10 (47.6%) | |
| Total Length of hospitalization | | | |
| Median (IQR) | 7.0 (5.0; 11.0) | 33.0 (27.0; 44.0) | < 0.001 |
| Type of hospitalization | | | |
| ARD (n (%)) | 0 (0.0%) | 21 (100.0%) | |
| ICU (n (%)) | 5 (14.3%) | 0 (0.0%) | - |
| Standard (n (%)) | 30 (85.7%) | 0 (0.0%) | |
| Length of hospitalization at ARO/ICU | | | |
| Days in ICU/ARD | ICU (n = 5) | ARO (n = 21) | |
| Median (IQR) | 10.0 (10.0; 13.0) | 30.0 (19.0; 38.0) | 0.010 |
| Oxygen therapy | | | |
| None | 14 (40.0%) | 0 (0.0%) | |
| Oxygen mask | 19 (54.3%) | 0 (0.0%) | |
| HFOT | 2 (5.7%) | 0 (0.0%) | - |
| NIV | 0 (0.0%) | 6 (28.6%) | |
| Mechanical ventilation | 0 (0.0%) | 15 (71.4%) | |
| High creatinine blood level | | | |
| Yes (n (%)) | 13 (37.1%) | 11 (52.4%) | 0.403 |
| No (n (%)) | 22 (62.9%) | 10 (47.6%) | |
| Abnormal liver function tests | | | |
| Yes (n (%)) | 17 (48.6%) | 18 (85.7%) | 0.013 |
| No (n (%)) | 18 (51.4%) | 3 (14.3%) | |
| Combination of high creatinine level + abnormal liver function tests | | | |
| Yes (n (%)) | 5 (14.3%) | 9 (42.9%) | 0.038 |
| No (n (%)) | 30 (85.7%) | 12 (57.1%) | |

*Continuous variables are presented as median (interquartile range - IQR). Categorical variables are presented as numbers (n) with percentages (%). Continuous variables for two groups were tested using the Wilcoxon test. Categorical variables were tested using the Chi-square test with Yates' continuity correction. C19 – COVID-19, ARD – Anesthesiology and resuscitation department, ICU – Intensive care unit.*

**Supplementary Table 2**: List of antibodies for FACS-phenotyping used in this study.

| Marker | Fluorochrome | Clone | Vendor |
|---|---|---|---|
| **CD10** | BV650 | HI10a | BD Horizon |
| **CD101** | R718 | V7.1 | BD Optibuild |
| **CD11b** | R718 | ICRF44 | BD Optibuild |
| **CD127** | APC ef780 | eBioRDR5 | Invitrogen |
| **CD14** | FITC | M5E2 | BD Pharmingen |
| **CD14** | BV510 | M5E2 | Biolegend |
| **CD141** | BV711 | 1A4 | BD Horizon |
| **CD142** | BV421 | HTF-1 | BD Optibuild |
| **CD16** | PE Cy5 | 3G8 | Biolegend |
| **CD172a** | FITC | 15-414 | Biolegend |
| **CD25** | APC | M-A251 | Biolegend |
| **CD3** | BV650 | UCHT1 | BD Horizon |
| **CD3** | PE Cy5.5 | SK7 | Invitrogen |
| **CD3** | APC Cy7 | OKT3 | Biolegend |
| **CD33** | BV711 | WM53 | SONY |
| **CD36** | PE Cy7 | 5-271 | Biolegend |
| **CD39** | BV605 | A1 | BD Horizon |
| **CD4** | BV510 | SK3 | Biolegend |
| **CD4** | BV650 | SK3 | BD Horizon |
| **CD45** | PerCP-Cy5.5 | HI30 | BD Pharmingen |
| **CD45RA** | PE Cy5.5 | MEM-56 | Invitrogen |
| **CD56** | PE Dazzle 594 | 5.1H11 | Biolegend |
| **CD56** | APC ef780 | TULY56 | Invitrogen |
| **CD56** | BV785 | 51.H11 | Biolegend |
| **CD57** | PE Cy7 | HNK-1 | Biolegend |
| **CD64** | BV605 | 10.led | BD Horizon |
| **CD66b** | PE | G10F5 | Invitrogen |
| **CD66b** | PE Dazzle 594 | G10F5 | Biolegend |
| **CD68** | PE Cy7 | Y1/82A | Biolegend |
| **CD8** | PE Cy5.5 | 3B5 | Invitrogen |
| **CD8** | FITC | SK1 | Biolegend |
| **CD85d** | AF647 | 287219 | BD Pharmingen |
| **CD86** | BV786 | IT2.2 | Biolegend |
| **CD89** | R718 | A59 | BD Optibuild |
| **CD95** | BV711 | DX2 | BD Horizon |
| **GITR** | BV421 | V27-580 | BD Horizon |
| **HLA-DR** | BV421 | L243 | BD Horizon |
| **PD-1** | BV605 | EH12.2H7 | Biolegend |

| | | | |
|---|---|---|---|
| **RAGE** | AF647 | A-9 | Santa Cruz Biotechnology |
| **TIGIT** | PE | TgMab-2 | BD Pharmingen |
| **TLR4** | PE | HTA125 | Invitrogen |
| **TREM-1** | BV786 | 6B1 | BD Optibuild |

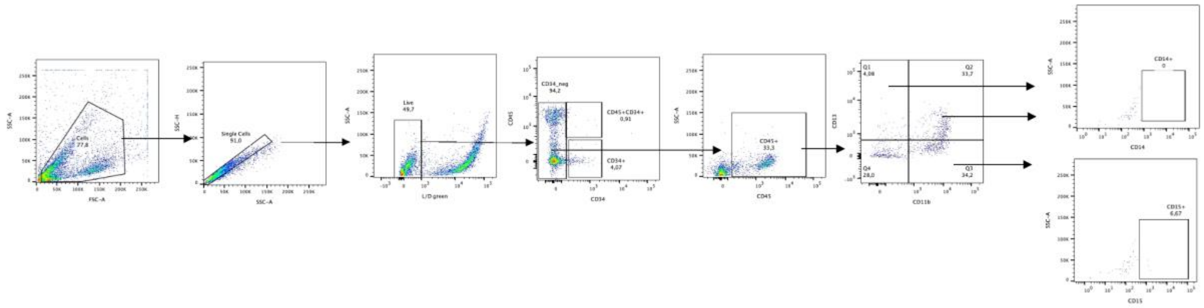


**Supplementary Figure 1**: Representative gating strategy for CFU phenotyping using FACS.

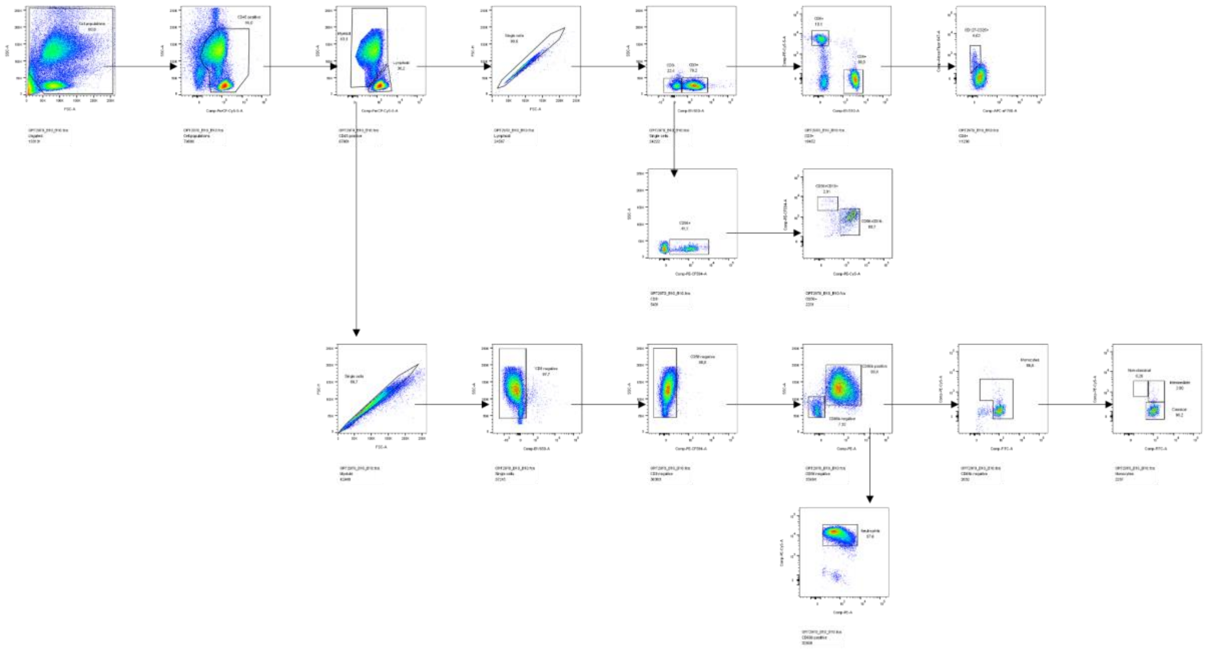

**Supplementary Figure 2:** Representative gating strategy to identify main immune cell populations in whole blood.

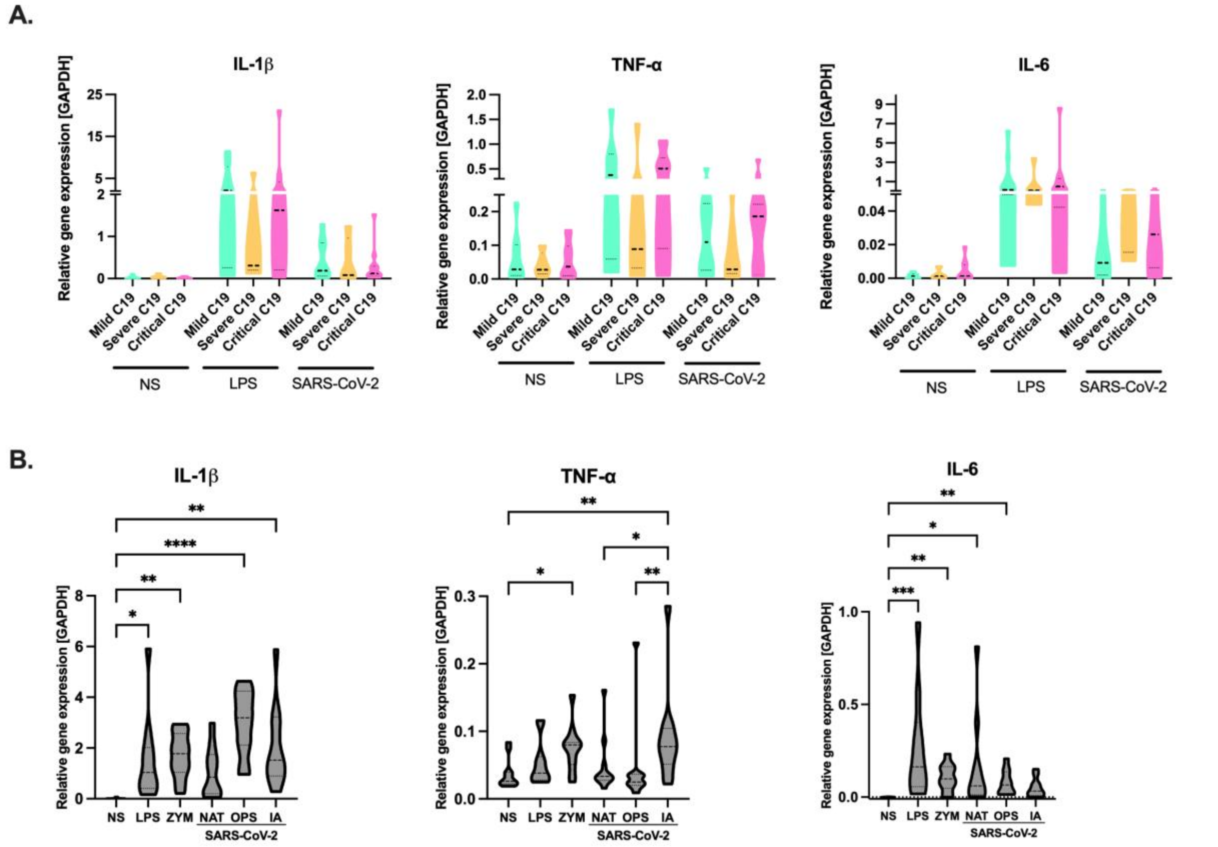


**Supplementary Figure 3: Response of monocytes isolated from post mild-to-critical COVID-19 and healthy volunteers to microbial challenge.**

Pro-inflammatory gene expression was measured at mRNA level using q PCR. A.) COVID-19 severity-associated response of monocytes to stimulation with LPS and SARS-CoV-2. B.) Monocyte responses to various microbial stimuli in healthy volunteers, regardless of their BCG vaccination status. Data were tested using Kruskal Wallis + Dunn's multiple comparison test. Statistically significant differences are indicated as follows: *P < 0.05, **P < 0.01, ***P < 0.001.

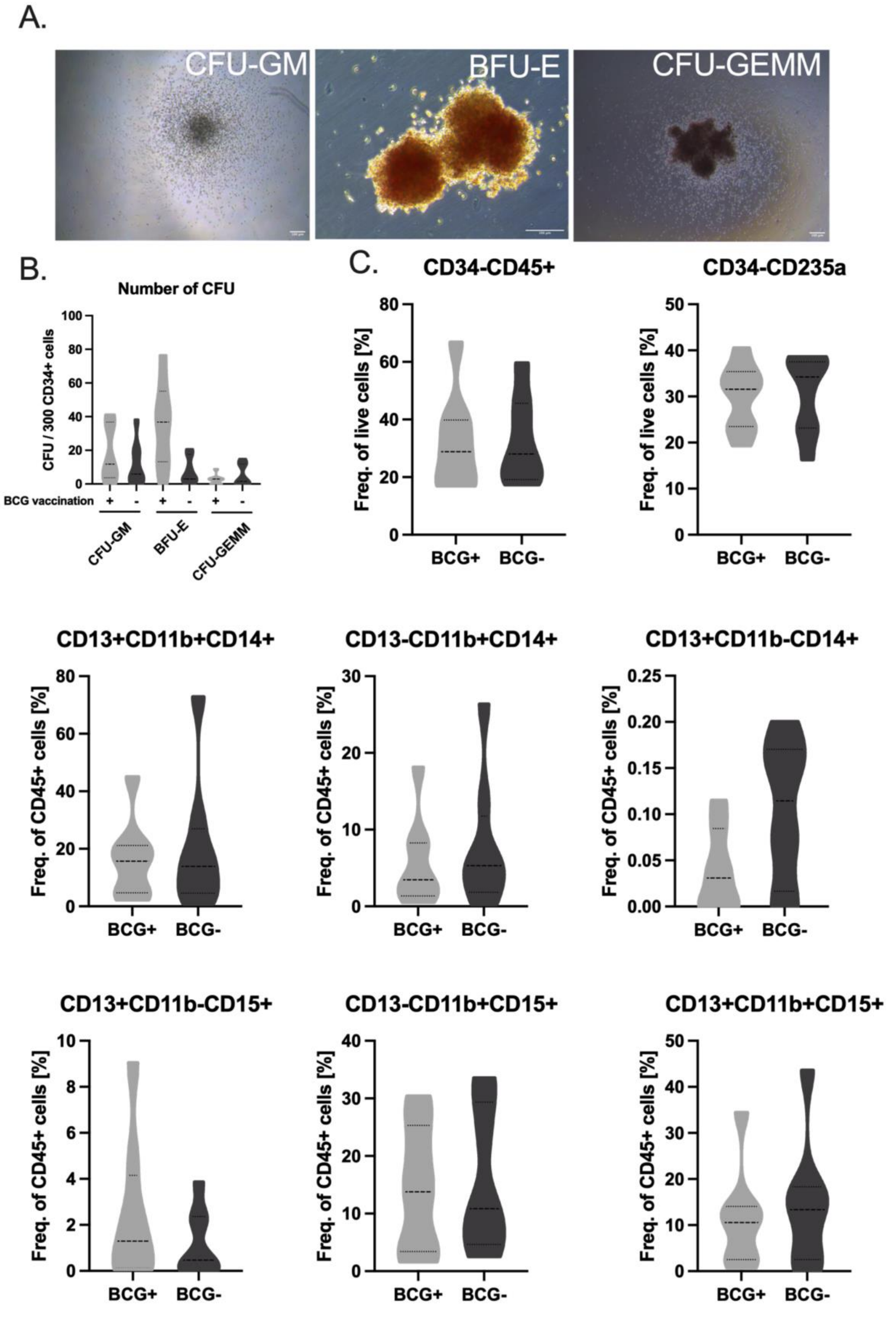


**Supplementary Figure 4: Methylcellulose colony forming assay.** A.) Representative images of colony formation in MethoCult ® media following 21-days of culture. B.) Total number

of CFU formed in MethoCult media. C.) FACS phenotyping of total CFU colonies. Statistical significance was tested using Kruskal Wallis followed by Dunn's multiple comparison test.
*BFU-E, burst-forming unit-erythroid; CFU-GM, colony-forming unit-granulocyte/macrophage; CFU-GEMM - multi lineage progenitors (granulocytes, erythrocytes, monocytes, and macrophages).*
